# Single-Cell, Human Sperm Transcriptomes and Variants from Fathers of Autistic and Healthy Children

**DOI:** 10.1101/640607

**Authors:** Delia Tomoiaga, Vanessa Aguiar-Pulido, Shristi Shrestha, Paul Feinstein, Shawn E. Levy, Jeffrey A. Rosenfeld, Christopher E. Mason

**Author notes:** Co-first authors (Delia Tomoiaga, Vanessa Aguiar-Pulido).

## Abstract

The human sperm is one of the smallest cells in the body, but also one of the most important, as it serves as the entire paternal genetic contribution to a child. This is especially relevant for diseases such as Autism Spectrum Disorders (ASD), which have been correlated with advance paternal age. Historically, most studies of sperm have focused on the assessment of a bulk sperm, wherein millions of individual sperm are present and only high-frequency variants can be detected. Using 10X Chromium single cell sequencing technology, we have assessed the RNA from >65,000 single sperm cells across 6 donors (scsperm-RNA-seq), including two of whom have autistic children and four that do not. Using multiple RNA-seq methods for differential expression and variant analysis, we found clusters of sperm mutations in each donor that are indicative of the sperm being produced by different stem cell pools. Moreover, by comparing the two groups, we have found expression changes that can separate out the two sets of donors. Finally, through our novel variant calling from single-cell RNA-seq methods, we have shown that we can detect mutation rates in sperm from ASD donors that is distinct from the controls, highlighting this method as a new means to characterize ASD risk.

## Introduction

Single-cell RNA sequencing (scRNA-seq) allows for the discovery and investigation of cellular subtypes. To date, this technique has not been employed on human germline tissues such as ova or sperm. Mature, motile sperm (spermatozoa) (**Fig. 1A**), are a challenging cell type to perform single-cell RNA sequencing on, and differ from typical somatic cells in several aspects. First, they are transcriptionally restricted cells, retaining only a small quantity of RNA per cell (∼50 fg) which exists in a fragmented or partially degraded state (Johnson et al. 2011; Mao et al. 2013; Sendler et al. 2013). Second, transcription ceases during the spermatid stage of spermiogenesis and sequential displacement of histones by transition proteins and eventually protamines (PRM1 and PRM2) takes place, along with nuclear remodeling (**Fig. 1B**) (Kistler et al. 1996). Third, spermatozoa exhibit a compact nucleus, minimal cytoplasm, a head-housed acrosome and a mitochondria-heavy midpiece, plus a long tail of ∼50um (**Fig. 1A**). This specific cell morphology paired with its ability to move rapidly, can also prove challenging for capturing of single-sperm, especially with microfluidic devices. These features taken as a whole make sequencing of single spermatozoa RNA more challenging, but also create an ideal paradigm for investigating transcriptome composition of sperm at a mature stage where new RNAs are not being produced and the ones retained may be of functional importance to the oocyte.

**Figure 1.**
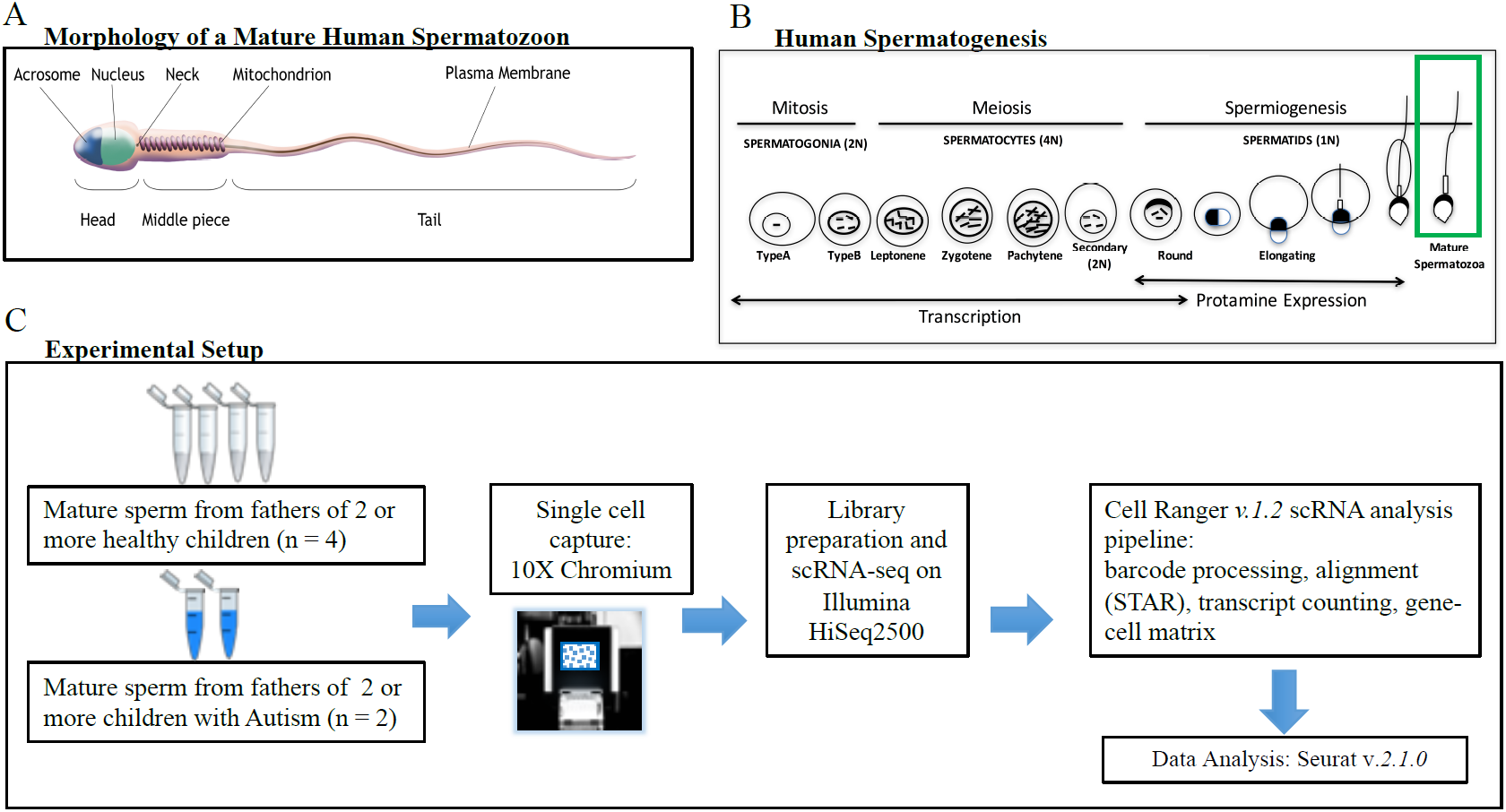
Overview of the single-cell RNA-seq experimental setup and background. (A) Morphology of mature human sperm. (B) The process of spermatogenesis and stages associated with transcription and expression. The consensus is that mature spermatozoa do not actively express RNA but that the RNA detected at this stage is retained from earlier stages. (C) Single-cell RNAseq human sperm experimental setup. Mature, motile human sperm fraction from two cohorts, i.e., males who fathered children with autism and males who fathered healthy, non-Autistic children, was loaded onto a Chromium 10X Genomics single-cell capture unit, library prepped and sequenced. Cell Ranger pipeline was used to calculate the expression matrix. Downstream data analysis was performed with Seurat.

The functions of the majority of RNAs in sperm remain unknown (Fischer et al. 2012). However, there has been recent evidence that spermatozoa may have a role in the regulation of early embryonic development by delivering functional RNAs to the oocyte during fertilization (Jodar et al. 2013; Krawetz 2005; Krawetz et al. 2011). The retained histones associate with telomeric sequences and shown to be the first sperm structures to respond to oocyte signals (Zalenskaya et al. 2000). Using the 10X Genomics Chromium platform (**Fig. 1C**), we performed single-cell RNA sequencing of six donors’ sperm samples obtained from a sperm bank with IRB approval (see methods). To explore putative differences in transcriptomes between spermatozoa from fathers of healthy offspring (Control) and spermatozoa from fathers of children diagnosed with Autism Spectrum Disorder (ASD), we used samples from both of these cohorts.

ASD is a complex neurodevelopmental condition with an often undetermined etiology (Sanders et al., 2011). It is classified as a Paternal Age Effect (PAE) disorder, since increased paternal age is associated with higher ASD risk. (Alter et al. 2011). Typically, two to three new mutations arise in sperm germ cells with each year of the father’s age (Francioli et al. 2015; Rahbari et al. 2016). Although hundreds of mutations increasing the risk of Autism have been identified, many cases have an unknown genetic component, thus limiting diagnostic breadth (Ansel et al. 2016). Here, we used a unique approach in performing Single Nucleotide Variant (SNV) calling on our RNA-seq data at single-cell resolution to uncover variants occurring in the sperm of affected donors (confirmed ASD children) and also performed a differential gene expression analysis between the two donor groups. Our data represent the first utilization of single-cell RNA sequencing and mutation detection at single-cell resolution in spermatozoa for identifying patterns of paternal transmission of disease. As such, this framework is the first step towards an RNAseq-based biomarker test in sperm for prediction of ASD risk and could lead to discoveries for other neurodevelopmental or PAE transmitted disorders as well.

## Results

### Sequencing results and filtering parameters

Among the six sperm samples, the number of barcodes with at least one gene varied between 72,507 in sample Healthy 3 and 97,064 in Autistic sample 2 (ASD2), while the number of genes detected varied between 18,238 in Healthy 4 (H4) and 21,701 in sample Autistic 1 (ASD1) (**Fig 2A**). For the downstream analysis, we excluded cells under the 25 UMI/cell threshold (**Fig 2B**), cells with fewer than 10 detected genes, and genes detected in less than 20 cells and cells with mitochondrial genes exceeding 40% of the total, this latter figure based on the fact that typical mature sperm contains about 40% mitochondrial content (Kistler et al. 1996). We detected 4,872 common genes among the 4 Healthy samples and 6,260 common genes between the 2 Autistic samples (**Fig. 2A**). Across all cells from all samples in the filtered dataset, which passed the above filtering thresholds, 4,266 genes remained (**Fig. 2A,2B**). Notably, after filtering with the above steps, the ASD samples showed a significant reduction (**Fig. 2C**) in the proportion of mitochondrial DNA (2.2×10^-16^, K-S test). We then used a normalization method with Seurat2 (Butler and Satija, 2017) to sub-sample the same number of read distributions across the common genes to compare the two groups. This led to comparable post-alignment distributions for the Control and ASH samples (**Fig. 2D-E**). After alignment, the number of cells in each sample were reduced (**Fig. 2F**) and the number of genes to 1,833, 4,239, and 632 in the ASD-specific, shared, and control-specific, respectively.

**Figure 2.**
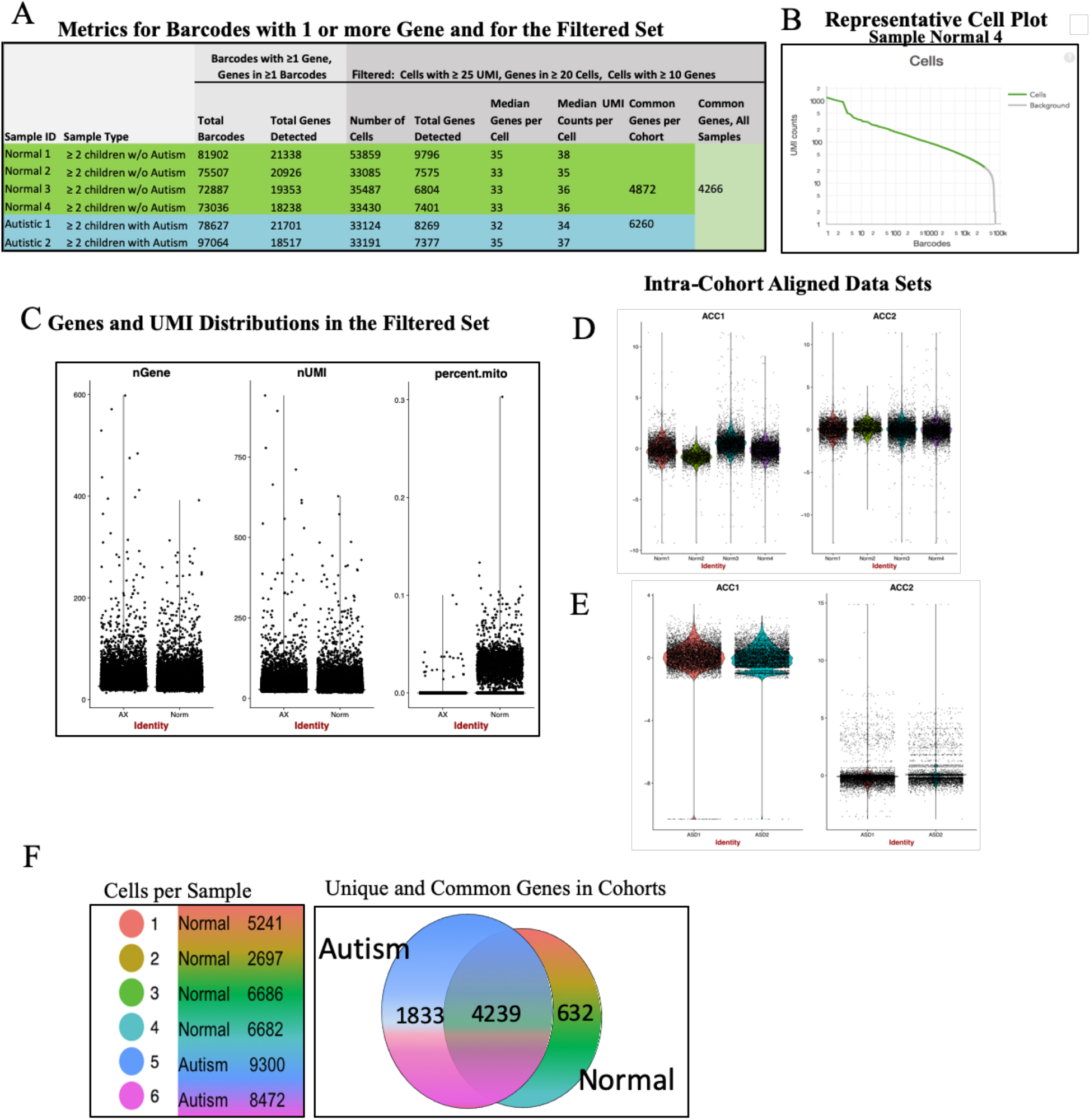
scsperm-RNA-seq profiles and metrics. (A) Metrics for i) barcodes containing 1 or more genes and ii)for the filtered set used in downstream analysis that includes all cells with 10 of more genes, all genes present in at least 20 cells, and all cells with 25 UMI or more (B) Representative cell plot showing the raw distribution of UMIs/barcodes for one healthy sample (C)The distribution of genes and unique molecular identifiers in the 6 datasets in the filtered set used for analysis (D), (E) Violin plots of the cohort-specific datasets post-alignment (F)

The highest transcripts in both healthy samples and ASD samples that were present in more than 40,000 cells included: PRM1, PRM2, TSSK6, DNAJC4, NUPR2, CRISP2 and SMCP (Table 1). Protamines replace about 85% of the histones in human sperm during maturation (Wykes and Krawetz 2003) and (as expected) are the most-expressed transcripts in our samples, appearing both in the highest number of cells and at the top of the gene-specific average expression values in our samples, confirming that we sequence mature spermatozoa and contamination from more immature precursor cells is absent or minimal. These are transcripts appear to be highly-specific to sperm, since they have lower or no expression in Human Universal Reference (UHUR) controls. (SEQC Consortium, 2014, Li et al, 2014).

We next examined the differences between the two cohorts. Overall, the gene expression signatures between the two groups was highly correlated (R^2^=0.86) and both groups contained signature genes for sperm and some differences (**Fig. 3A**). Specifically, we observed high expression of CCDC91 and AKAP1 in both cohorts (**Fig. 3B**), which is important since CCDC91 is a marker of mature motile sperm and AKAP1 preserves mitochondrial integrity. Differential expression analysis comparing the cells in the Autism and the Healthy samples revealed 2,114 differentially-expressed genes (DEGs), of which 1247 were increased and 867 were decreased in the ASD samples (p-value, *Bonferroni*-adjusted for multiple hypothesis testing < 0.05). When the minimum threshold for marker expression was set such that a transcript was present in both of the two groups and >1% percent of cells, the DEG list was reduced to 687 differentially expressed genes between the ASD and the Control groups, with 348 genes showing an increase and 339 transcripts showing a decrease in the ASD set.

**Figure 3.**
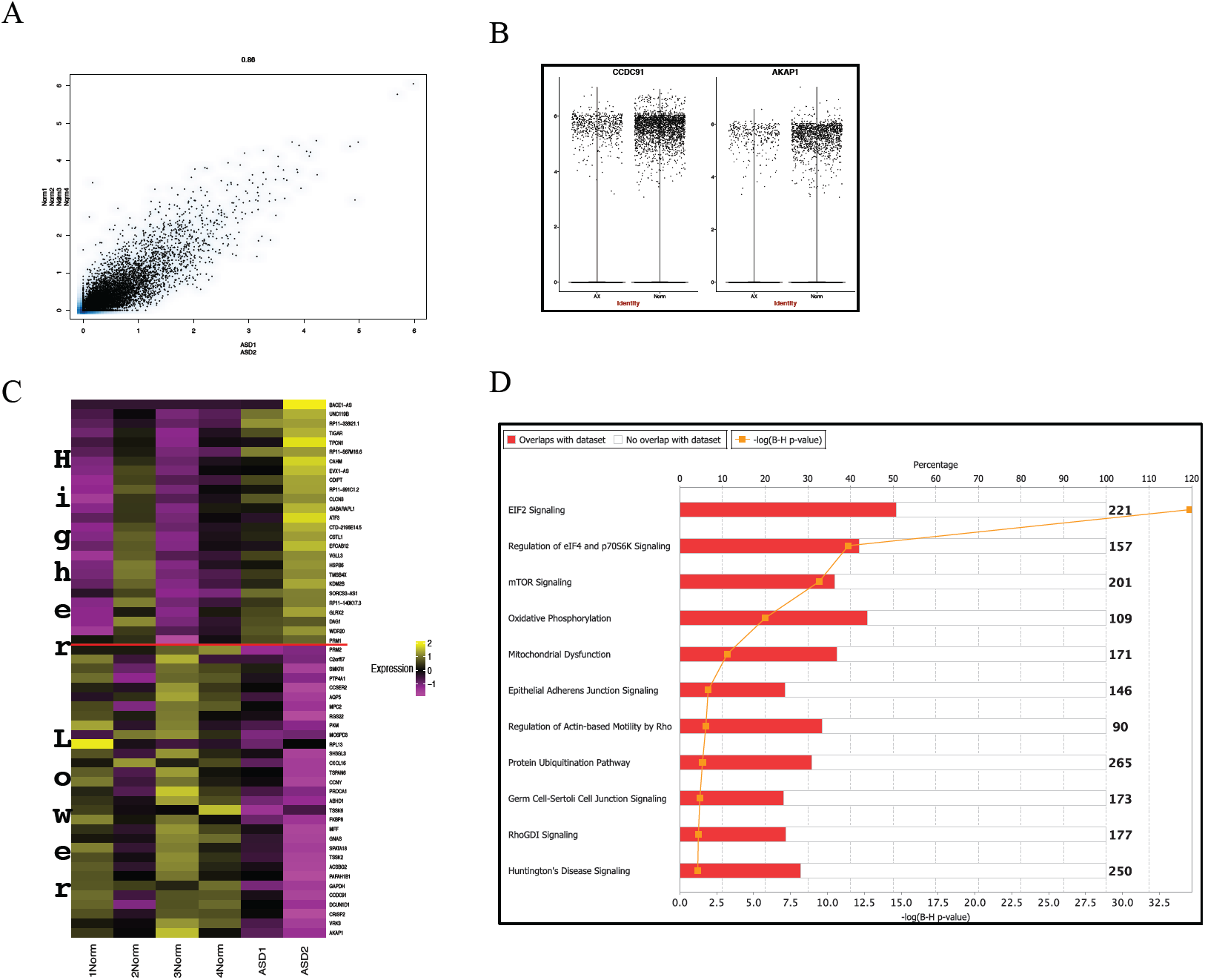
Distinctive profiles between cohorts. (A) Heatmap of DEGs between the Normal and the Autism cohort. Each rectangle represents the scaled average expression of the single cells for a specific gene. (B) Pathways enriched in DEGs, the red bar represents the ratio of DEGs in the pathway to total genes. (E) CCDC91 is marker of mature motile sperm and is higher in Normal. AKAP1 preserves mitochondrial integrity. (F) Scatter plot comparing the range in scaled average expression in each cohort. Each dot represents a gene expression value across all single-cells.

We then examined these DEGs for specific gene and pathway enrichments (**Fig. 3C, 3D**). Using the Qiagen Ingenuity Pathway Analysis tool, we observed several pathways significantly enriched (-log of p-value 1.54 to 34.6), including: EIF2 signaling, regulation of eILF4 and p70S6K signaling, mTOR signaling, oxidative phosphorylation, mitochondrial dysfunction, epithelial adherens junction signaling. (**Fig. 3D**). These pathways are all involved in sperm cell function and processing, and link to known neurodevelopmental pathways.

To explore the mutational landscapes of the individual sperm cells, we developed a new method for calling variants in scsperm-RNA-seq data (see methods). First, we examined the distribution of variations across the cells. For each sample, there were between 194 and 302 SNPs present, with non-synonymous SNPS found in genes including CARHSP1, CRISP2, DNAJC4, NUPR2, PRM1, PRM2, SMCP. To increase the stringency and to remove false positives, we further filtered the data to only include variants that were found in at least 100 sperm cells. At this level, we only found non-synonymous sperm in the PRM1 and PRM2 genes (**Fig. 4A, 4B**). Notably, we were able to discern distinct mutations in the sperm with enough coverage, which is, to our knowledge, the first detection of such variants from individual sperm RNA.

**Figure 4.**
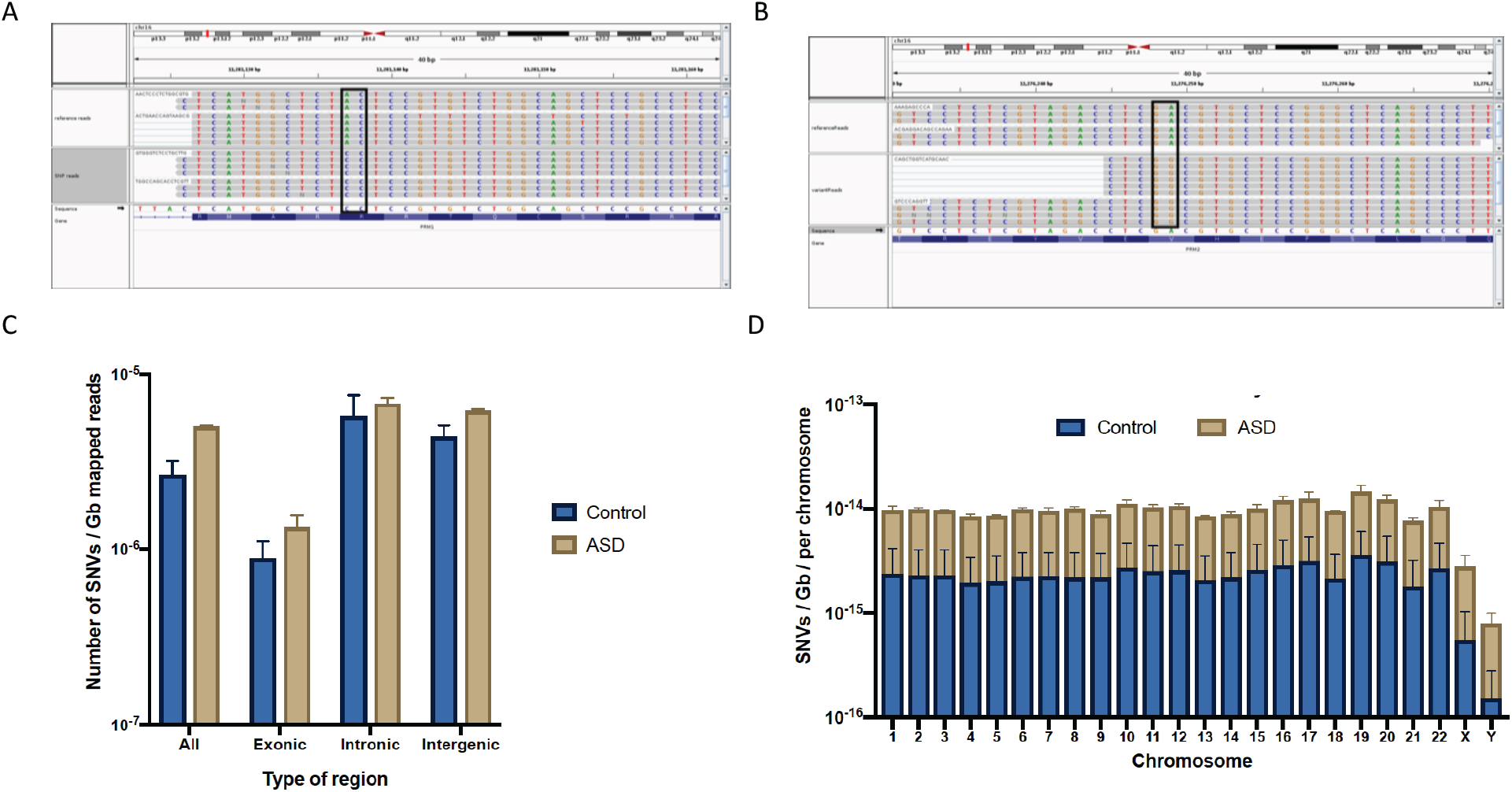
SNVs from single sperm cells. Using the Integrative Genome Viewer (IGV), data is shown from alignments to the reference genome from single cells. (A) PRM1 variants are shown as coverage tracks (rows) with the reference genome on top and the donor variants on the bottom row. (B) Same as A, but shown for the same donor in PRM2. (C) Number of SNVs with an allele frequency < 0.001 in gnomAD per chromosome, normalized to the size of the read (bp) multiplied by the total number of reads mapped to the transcriptome. (Values are shown in a log10 scale). (D) SNVs distribution across chromosomes.

Given recent reports of using 10xGenomics single-cell RNA-seq data for calling variants within genes and nearby regions (Allegra et al., 2019), we next examined the overall distribution of variants in the ASD samples and the controls. By using the GATK RNA-seq pipeline for variant calling and annotation (see methods), we compared our called variants to those annotated as rare SNVs (allele frequency <0.001) within the public genome aggregation (gnoMAD) database (https://gnomad.broadinstitute.org/) (Karczewski et al., 2019). These data indicated that the total number of variants found in the ASD samples was higher overall (**Fig. 4C**), as well as in total exonic, intronic, and intergenetic regions. This trend held true across both autosomes and sex chromosomes (**Fig. 4D**), and indicated a new metric of expressed, rare variants from sperm samples that stratifies based upon ASD phenotype.

## Discussion

These results demonstrate that scsperm-RNA-seq is a promising method to profile gene expression and mutational dynamics of the transcriptomes of individual sperm cells. As expected, the highly expressed genes in both the ASD and Control samples displayed an overlap with genes in pathways for sperm maturation, early embryonic development, cell growth, and proliferation. Although the DEG and pathway differences observed between the cohorts can provide potential leads or serve as biomarkers for sperm health or function, they will need to be validated in a larger donor cohort size.

Nonetheless, the differences between the cohorts reveal distinct expression landscapes and pathways, such as enrichments for mTOR signaling and eIF2 regulation in the ASD samples. The mTOR signaling pathway has recently been highlighted as a potential target of Autism (Sato, 2016), while the eIF2 pathway is involved in the inhibition of CREB, a transcription factor required for long-lasting synaptic plasticity and long-term memory and has recently been investigated in connection with several neurodegenerative disorders (Trinh and Klann 2013). Also, RPL10 was the most decreased transcript in our list of DEGs in the ASD cohort (log_2_ decrease of 1.98), and mutations in RPL10 (e.g c.*190C>G, c.453C>A) can lead to gene dysfunction and confer a risk of X-linked autism in humans. These specific mutations have not been identified in our data, but a similar ribosomopathy may be caused by the lack of RPL10 RNA expression in the Autistic sperm samples.

Also, several genes with decreased expression in the ASD vs. Normal samples have documented, central roles in sperm development. PACRG is recognized to increase both spermiogenesis and spermatogenesis, OVOL1 and DDX3Y both increase the rate of spermatogenesis, TPP2 and CLU affect the maturation of sperm. Moreover, AGFG1 increases the rate of acrosome formation and ACE increases binding of sperm. For several genes displaying increased expression in the ASD samples in our DEG analysis, studies exist that show correlation with an increase in the ASD phenotype: DRD3, CADPS2, TCF7L2 (Correia et al. 2010). Given sufficient depth from sequencing, we have also shown that an investigation of sequence variants in RNA from sperm is also possible. These variants will be limited to the genes that are highly expressed in the sperm and in the 3’ ends of the sequenced RNAs.

Work in a larger replication cohort of donors based on our methods could further help to isolate patterns useful for understanding paternal transmission of ASD and may help provide missing link between genome and phenotype, not only for individual sperm but across the distribution of all sperm cells. Such methods could, in principle, eventually be used as a screen of sperm to help determine which males are at a higher risk of having a child affected with Autism or another PAE disorder, and also provide a high-resolution map of sperm health.

## Methods

### Samples

Sperm samples were purchased from California Cryobank (Los Angeles, CA). All of the donors signed an informed consent with the cryobank agreeing to the use of their samples in research. Since the donor sperm samples at the cryobank are purchased by women to use for insemination, many of the donors have a large number of children. Interestingly, there are cases where a donor will have one of his many children having Autism with the rest being unaffected, indicating that there are other causes of autism besides paternal genetics. For this reason, we required that our Autism donors have evidence of at least 2 children with Autism. The control donors needed to have multiple children with no evidence of Autism. The samples were of IUI-quality, purified by the cryobank with the standard protocol for enrichment of mature, motile spermatozoa, and were shipped on liquid nitrogen from the cryobank to the lab where they were thawed for analysis. Their donation and use was approved by the Cryobank IRB.

### Cell preparation

Frozen sperm cell vials were obtained from California Cryobank Biotech. Sperm cells were provided with expected post-thaw cell count. Frozen cells were rapidly thawed in a 37°C water bath. Thawed cells were centrifuged at 300 rcf for 10 minutes and washed twice with 1x PBS containing 0.04% bovine serum albumin, then resuspended in PBS at room temperature.

### Single cell library construction and sequencing

Cell suspension post-washing was loaded on the 10X Chromium System (10x Genomics, Pleasanton, CA) for single cell isolation into Gel Bead Emulsions (GEMs) as per manufacturer’s instruction in Chromium Single Cell 3’ Reagent Kits v2 User Guide, Rev A (Zheng et al., 2017) using Chromium Single Cell 3’ Solution (Chromium Single Cell 3’ Chip Kit v2 [PN-120236], Gel Bead kit v2 [PN-120235]. The input cells per channel in the chip were targeted around ∼1 million cells based on provided initial post-thaw cell count from California Cryobank Biotech. However, the loss of cells during washing would reduce the actual cell counts to input significantly less than targeted cells.

Sperm samples that successfully generated proper GEMS were further processed for GEM-RT incubation, cDNA amplification and subsequent single cell library construction using Chromium™ Single Cell 3’ library Kit v2 [PN-120234] following the manufacturer’s protocol. Barcoded final libraries were quantified by Qubit^®^ 2.0 Fluorometer (Invitrogen) and qPCR (KAPA Biosystems Library Quantification kit) and fragment size profiles were assessed from Agilent 2100 BioAnalyzer. All libraries were sequenced on Illumina Hiseq 2500 with 2X 100 paired-end kits using following read length: 26 bp Read 1, 8 bp i7 Index and 98 bp Read 2.

Cellranger (*v 1.2*) single cell pipeline (https://support.10xgenomics.com/single-cell-gene-expression/software/overview/welcome) was used for demultiplexing libraries, using cellranger mkfastq to generate FASTQ files. STAR alignment, barcode/UMI processing and counting were conducted by the Cellranger count pipeline. Barcode, UMI and duplicate sorting are further described (Zheng et al., 2017).

### Data Analysis

For the data analysis, we excluded cells under the 25 UMI/cell threshold, cells with fewer than 10 detected genes, and genes detected in less than 20 cells. Cells with mitochondrial genes comprising more than 40%, a typical mitochondria fraction in a healthy spermatozoon (ref), were excluded. Multiple pairwise alignment to remove batch effects and allow for integrated analysis was performed on all samples for each cohort as described and implemented in R toolkit Seurat (*Seurat v.2.1.0 and 2.2.0)*(Butler and Satija 2017). Briefly, the Autism and the Healthy control filtered datasets were randomly subsampled to 12,000 cells per sample, including 24,000 cells total for Autistic and 48,000 cells for the Control dataset. The cells were subsampled based on common genes, the expression normalized, scaled and aligned into a conserved low-dimensional space using non-linear warping algorithms. Canonical correlation vectors are aligned to normalize for differences in feature scale, an approach robust to shifts in population density. Variable genes were detected across the data sets. The aligned merged datasets are used for downstream analysis. The Autism dataset (ASD) includes two samples, ASD1 with 9300 cells and ASD2 with 8472 cells. The healthy control dataset (Healthy) contains 4 samples: Healthy 1-4, each containing 5241 cells, 2697 cells, 6686 cells, and 6682 cells, respectively.

We performed differentially-expressed gene (DEG) analysis between the all the single cells in the aligned cohort-specific datasets. For detecting differentially expressed genes between the sperm samples from fathers of children with Autism and those of healthy children, we performed a Wilcoxon rank sum test at single-cell level. A t-distributed stochastic neighbor embedding (t-SNE) analysis to visualize cells in a two-dimensional space was done. Based on the relative position of the cells on the t-SNE plot, unsupervised graph-based clustering was performed with the *FindClusters* function in Seurat and unique cluster marker genes were identified.

#### Variant calling

The BAM file produced by the CellRanger software is not designed to easily allow for variant calling and needs to be modified. In particular, the reads from each individual cell are marked with the CB tag in the BAM file whereas a typical BAM file would record each set of reads from the same source as a read-group with the RG tag. We developed a cleaning algorithm (NNN.pl) that converted the BAM file and also removed any of the cells that had less than 100 reads per cell. We then used the Freebayes variant caller on the converted BAM files and we removed any variants with a quality of <20 and required that a SNP be present in at least 10 individual single cells. Thus, our SNPs were relatively rare in the sperm population, but were not only found in an individual cell.

#### Code for variant pruning

~~~
## convertBAMfull.perl
use strict;
my($inLine,$rgTag,$cbTag,$rawTag,%allRG,$thisOut,$thisRG,@inSplit,$fixedLine,$i,$counter
,%allReadLines,%rgCounter);

keys(%allReadLines) = 100000000;
keys(%rgCounter) = 10000000;
keys(%allRG) = 10000000;
open(INTAKE,$ARGV[0]);
while($inLine=<INTAKE>) {
  $counter ++;

  if($counter % 100000 ==0)
  {print STDERR “$counter\n”;
  }

  if($inLine =∼/^@/) {# a header line
      print $inLine;
      next;
  }

  @inSplit=split(/\t/,$inLine);
  $cbTag= $inSplit[18];
  $rawTag= (split(/:|-/,$cbTag))[2];
  $rgTag = “RG:Z:$rawTag”;

  $allRG{$rawTag}=$rawTag;

  $inSplit[18]=$rgTag;

  $fixedLine=join(“\t”,@inSplit);
  push(@{ $allReadLines {$rawTag}}, $fixedLine);
  $rgCounter{$rawTag} ++;
}

  foreach $thisRG(keys(%allRG)) { #print tags with enough reads
     if($rgCounter{$thisRG} > 5) {
   print “\@RG\tID:$thisRG\tSM:$thisRG\n”;
   }
}
foreach $thisRG(keys(%allRG)) { #print tags with enough reads
   if($rgCounter{$thisRG} > 5) {

   foreach $thisOut (@{$allReadLines{$thisRG}}) {
     print $thisOut;
     }
   }
}
~~~

#### SNV calling and annotation in bulk

Genetic variants were called using the Broad Institute’s GATK [The Genome Analysis Toolkit: a MapReduce framework for analyzing next-generation DNA sequencing data McKenna A, Hanna M, Banks E, Sivachenko A, Cibulskis K, Kernytsky A, Garimella K, Altshuler D, Gabriel S, Daly M, DePristo MA, 2010 GENOME RESEARCH 20:1297-303] Best Practices for RNA-seq variant calling. Duplicates were marked and aligned reads were sorted using Picard tools. The SplitNCigarReads was used to split reads into exon segments and to clip reads overhanging intron regions. Variants were called using the HaplotypeCaller and single nucleotide polymorphisms (SNPs) were extracted using SelectVariants. Hard filtering was carried out using VariantFiltration to remove artifacts due to clusters of at least 3 SNPs in windows of 35bp, as recommended. Finally, variants with a coverage <20X for the alternative to the reference genome were not included in the analysis. The remaining variants were annotated using the most updated version (96) of Ensembl Variant Effect Predictor (VEP) [McLaren W, Gil L, Hunt SE, Riat HS, Ritchie GR, Thormann A, Flicek P, Cunningham F. The Ensembl Variant Effect Predictor. Genome Biology Jun 6;17(1):122. (2016)].

*(Version if using the same denominators)*

#### SNP data analysis (all, coding, noncoding)

All variant counts were normalized for each sample using the following formula:

Normalized number of variants_sample_i_ = Number of variants_sample_i_ / (Reads confidently mapped to the transcriptome_sample_i_*Read size)

VEP filter was used to extract variants with an allele frequency in gnomAD < 0.001 or absent from the database. The total number of rare variants per chromosome was calculated and normalized following the subsequent formula:

Normalized number of rare variants_sample_i,chr_i_ = [Number of rare variants_sample_i,chr_i_ / (Reads confidently mapped to the transcriptome_sample_i_*Read size)] / Length of chr_i

## Acknowledgements

We would like to thank the Epigenomics Core Facility at Weill Cornell Medicine, as well as the Starr Cancer Consortium (I9-A9-071) and funding from the Irma T. Hirschl and Monique Weill-Caulier Charitable Trusts, Bert L and N Kuggie Vallee Foundation, the WorldQuant Foundation, The Pershing Square Sohn Cancer Research Alliance, NASA (NNX14AH50G, NNX17AB26G), the National Institutes of Health (R25EB020393, R01NS076465, R01AI125416, R01ES021006, 1R21AI129851, 1R01MH117406), TRISH (NNX16AO69A:0107, NNX16AO69A:0061), the Bill and Melinda Gates Foundation (OPP1151054), the Leukemia and Lymphoma Society (LLS) grants (LLS 9238-16, Mak, LLS-MCL-982, Chen-Kiang) and the Alfred P. Sloan Foundation (G-2015-13964).

